# Preexisting hippocampal network dynamics constrain optogenetically induced place fields

**DOI:** 10.1101/803577

**Authors:** Sam McKenzie, Roman Huszár, Daniel F. English, Kanghwan Kim, Euisik Yoon, György Buzsáki

## Abstract

Neuronal circuits face a fundamental tension between maintaining existing structure and changing to accommodate new information. Memory models often emphasize the need to encode novel patterns of neural activity imposed by “bottom-up” sensory drive. In such models, learning is achieved through synaptic alterations, a process which potentially interferes with previously stored knowledge ^1-3^. Alternatively, neuronal circuits generate and maintain a preconfigured stable dynamic, sometimes referred to as an attractor, manifold, or schema ^4-7^, with a large reservoir of patterns available for matching with novel experiences ^8-13^. Here, we show that incorporation of arbitrary signals is constrained by pre-existing circuit dynamics. We optogenetically stimulated small groups of hippocampal neurons as mice traversed a chosen segment of a linear track, mimicking the emergence of place fields ^1,14,15^, while simultaneously recording the activity of stimulated and non-stimulated neighboring cells. Stimulation of principal neurons in CA1, but less so CA3 or the dentate gyrus, induced persistent place field remapping. Novel place fields emerged in both stimulated and non-stimulated neurons, which could be predicted from sporadic firing in the new place field location and the temporal relationship to peer neurons prior to the optogenetic perturbation. Circuit modification was reflected by altered spike transmission between connected pyramidal cell – inhibitory interneuron pairs, which persisted during post-experience sleep. We hypothesize that optogenetic perturbation unmasked sub-threshold, pre-existing place fields^16,17^. Plasticity in recurrent/lateral inhibition may drive learning through rapid exploration of existing states.

---

The ability for hippocampal circuits to imprint a random, novel pattern was tested in transgenic mice in which channelrhodopsin2 (ChR2) was expressed in excitatory neurons (N = 6 CaMKIIα-Cre:: Ai32 mice). Stimulation was achieved through μLED illumination^18^, which was delivered as mice ran on a linear track (1.2 m) for water reward (Figure 1A). After ten baseline trials, stimulation (1s half sine wave) was given for one to ten trials (see Supplemental Table 1) at a fixed position and running direction that changed daily. Optogenetic stimulation induced highly focal drive in CA1 neurons (N = 715; rate change on stimulated shank, Wilcoxon signed-rank test p = 4.1^−183^, median number stimulated = 12 neurons, range 1-50 neurons), as pyramidal cells on the neighboring shanks (≥ 250μm away; N = 420) showed no increase in firing even at the highest stimulation intensity (Figure S1, rate change on non-stim shank, Wilcoxon signed-rank test p = 0.40, median number stimulated = 0 neurons, range 0-5 neurons). On the track, pyramidal cells on the non-stimulated shanks were moderately suppressed in the stimulation zone, as compared to baseline, non-stimulated trials (mean rate change −0.10 ± 0.07 Hz, sign-test, p =7.34^−5^). In CA3 and the dentate gyrus (DG), the activity of pyramidal neurons on neighboring shanks increased (Figure S1), as expected from the recurrent excitatory connections in these regions. Local inhibitory interneurons were also strongly driven (Figure S1), likely through local synaptic drive^18^. These results show that focal stimulation drove feedback inhibition in CA1, and feedback excitation and inhibition in CA3 and the dentate gyrus.

**Figure 1.**
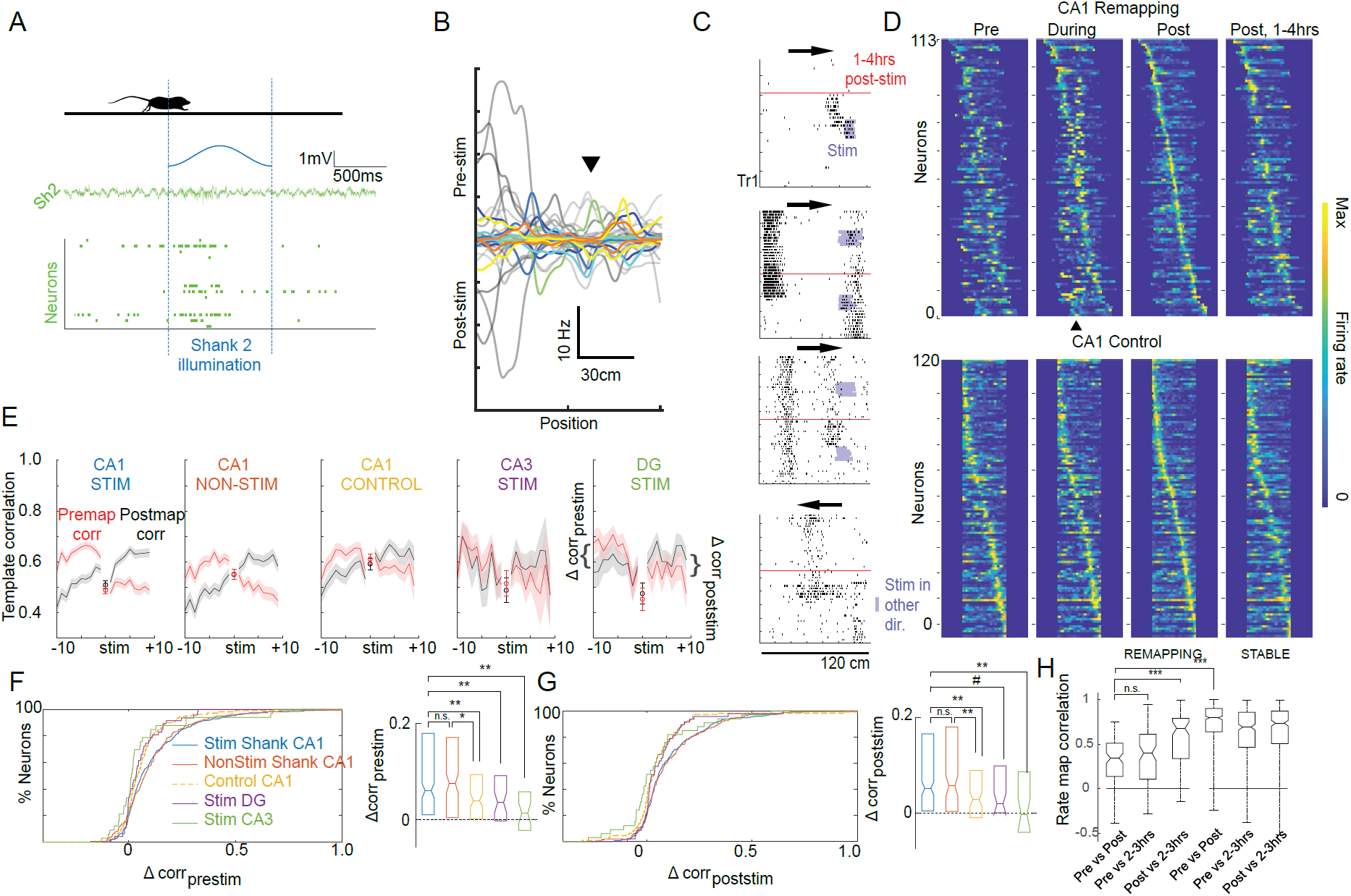
Focal optogenetic stimulation in CaMKIIα-Cre::Ai32 mice induced place field remapping in CA1, but not CA3 or DG. *(A)* Top 1s half-sine wave was delivered at a fixed position on a linear track (dashed line). Example raster and LFP showing drive on the stimulated shank (Sh2) *(B)* All place fields for an example stimulation session before (Pre-stim) and and after (Post-stim) optogenetic stimulation. ▴ marks stimulation location. Colored traces demark significantly remapping cells (see Methods). *(C)* Example CA1 cells that remapped after focal light delivery (light blue). Arrows show running direction. *(D) Top*, Population firing rate maps for remapping CA1 cells recorded on a stimulation session. Rate maps aligned to stimulation location (▴). Neurons sorted by the location of the peak firing post-stimulation (Post) and the rate normalized to the peak rate observed outside of stimulation. *Bottom*, Rate maps for all CA1 neurons recorded on Control days without stimulation (see Figure S2 for CA3 and DG) *(E)* Correlations of trial-by-trial rate maps against templates defined by pre-stimulation activity (Premap corr) and against post-stimulation activity (Postmap corr) *(F,G)* The difference between these templates (Δcorrprestim and Δcorrpoststim) shows the degree of place field remapping. *(H)* For remapping CA1 neurons, place field reorganization was stable 2-3 hours after stimulation. Neurons that did not remap with stimulation (Stable) maintained long-term place field stability. *** p < .001, ** p <.01, * p < .05, # p =0.1.

We next tested whether optogenetically-induced place fields^19,20^ persisted in the stimulation zone. In contrast to prior studies that manipulated single neurons only^1,14,15^, we found that novel place fields (see Supplemental Methods for place field criteria) emerged and disappeared both within and outside of the stimulation zone (Figure 1B-D, Figure S2C-E), and in both the stimulated and non-stimulated running direction (Figure S2D). To quantify this remapping, we adopted a template-based approach in which the trial-by-trial rate maps were correlated with the mean place field map prior to stimulation versus one derived from post-stimulation place fields. Large divergence in the Pre-vs Post-template matches is indicative of place field reorganization (Figure 1E). Stimulation caused place field remapping in area CA1 (N = 323 place cells on the stimulated shank) as compared to non-stimulated Control sessions (N = 120 place cells) conducted in the same mice on different days (Figure 1D-G; Pre-stim trials, median Δr Stim= 0.06 ± 0.01; Con = 0.04 ± 0.02, Mann-Whitney U-test p = 0.009; Post-stim trials, median Δr Stim= 0.05 ± 0.01; Con = 0.02 ± 0.02, Mann-Whitney U-test p = 0.006). Neurons with place fields on non-stimulated shanks (N = 160) also showed stronger place field remapping relative to Controls (Figure 1D-G, Figure S2; Pre-stim trials, median Δr Stim= 0.08 ± 0.01, Mann-Whitney U-test, p = 0.02; Post-stim trials, median Δr = 0.06 ± 0.01, Mann-Whitney U-test, p = 0.002). The remapping observed in neurons recorded in the CA1 region was stronger than that observed for neurons recorded in CA3 (Figure 1E-G; Figure S2; N = 33 place fields, Pre-stim trials, median Δr Stim= 0.01 ± 0.02; p = 0.001; Post-stim trials, median Δr = -0.003 ± 0.03; CA3stim vs CA1stim p = 0.007) and the dentate gyrus (Figure 1E-G; Figure S2; N = 47 place fields,; Pre-stim trials, Δr Stim = 0.03 ± 0.02; p = 0.02; Post-stim trials, median Δr = 0.02 ± 0.02; DGstim vs CA1stim p = 0.07), whose place field stability was less affected by focal optogenetic stimulation. These results show rapid plasticity in a region dominated by lateral inhibition (CA1) and relative stability in regions with denser recurrent connections (CA3 and the dentate gyrus).

For CA1 neurons that showed a large mismatch (Δr > 0.25) between the pre- and post-stimulation place field templates (i.e. those that remapped; N = 113 on both stimulated and non-stimulated shanks, out of 483 total; vs N = 6/120 in Controls; Two proportion Z-test = 4.03, p < 0.00001), rate within the pre-stimulation place field decreased as rate within the new fields increased (Figure S3A,B,D). The locations of the place fields post-stimulation were not clustered around the stimulation zone (Figure S3C), showing reorganization of the track representation in CA1 stimulation sessions. A second recording session revealed that the remapping in CA1 was stable after 1-4 hours (median = 11,253s, st.dev = 3,672s, min = 4,237s) in the homecage (Figure 1H). By definition, the correlation between pre- and post-stimulation place field templates was lower for cells that remapped (median corr_PREvsPOST1_= 0.34 ± 0.02) as compared to stable cells (median corr_PREvsPOST1_= 0.80 ± 0.01; Mann-Whitney U-test p = 6.3^−46^). Although the time between the first and last stimulation (median = 382.0s, st. dev = 178.0s, max = 783s) was shorter than the interval between the first and second recording session, cells that remapped showed post-stimulation place fields that were more similar to those recorded during the second recording session (median corr_Post1vsPOST2_= 0.68 ± 0.03) than to those place fields prior to stimulation (Mann-Whitney U-test corr_PREvsPOST1_ vs corr_POST1vsPOST2,_ p = 6.9^−11^), thus suggesting that drift in place field coding or systematic recording instability are unlikely to explain remapping. We also found no difference in spike sorting quality between the remapping and non-remapping sub-populations, as measured by L-ratio (p = 0.85) and isolation distance (Mann-Whitney U-test p = 0.24). These results show long-term stability of optogenetically induced remapping in CA1.

For CA1 neurons that showed large template differences (N = 113), or showed a novel place field emergence (see Supplemental Methods; N = 31 neurons), the induced fields could be predicted from their firing patterns even prior to stimulation (Figure 2A,B). The firing rate in the location in which a novel field would emerge was significantly higher than a chance distribution (circular shift of place field), suggesting the prior existence of sub-threshold place-specific drive (Wilcoxon signed-rank test p<10^−10^) at the new place field center ^11,17^. We hypothesized that, prior to remapping, neurons were already part of an assembly tuned to the future place field location. If such preexisting structure exists, we reasoned that when the remapping neuron fires, other simultaneously recorded neurons should predict that the mouse occupies the future place field location. To quantify this prediction, for each remapping cell, we calculated the spike-triggered Bayesian posterior-probability of occupying the new place field location, using spikes from all other pyramidal cells (Figure 2B). Prior to stimulation, spike-triggered decoding to the new field location was increased relative to random moments on the track (Figure 2C), showing a pre-existing fine-timescale coupling with co-tuned neurons. After stimulation, synchronization with co-tuned neurons increased, as quantified by enhanced decoding to the new field (Figure 2C). Thus, optogenetic activation unmasked a pre-existing place field and enhanced coincident activity of neurons co-tuned for that new location.

**Figure 2.**
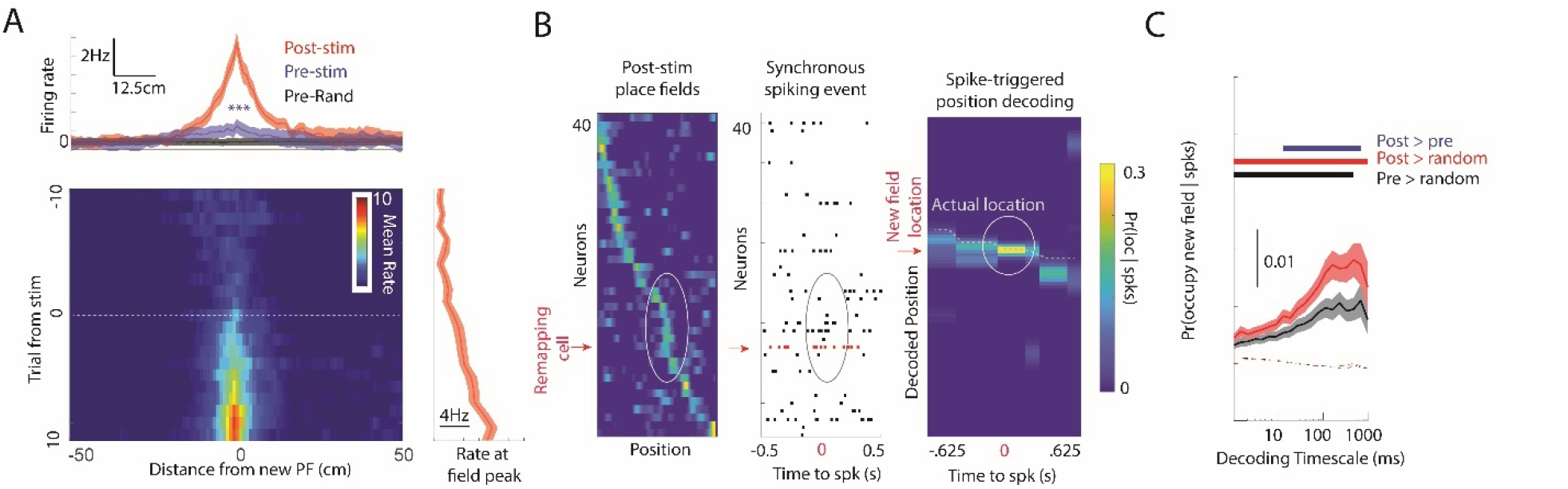
New fields emerge through strengthening of existing assembly membership. *(A)* For neurons that shifting (N = 113) or emerging (N = 31) place fields, firing in the future place field location was already higher than chance. *(B)* Strong Bayesian decoding to the post-stim place field center, Pr(occupy new field | spks), was used as a proxy fine-timescale synchrony with an ensemble of co-tuned neurons. *Left*, Example post-stimulation place fields sorted by peak location. Remapping neuron shown by red arrow. *Middle*, Spike-triggered raster centered on the spike of the remapping neuron showing synchrony (ellipse) with other neurons with similar place fields. *Right*, Posterior probability map showing high likelihood of occupying the new place field at the time of spiking. Dashed line: actual location of the mouse. *(C) Top* After stimulation, the spike triggered Pr(occupy new field | spks) increased, and was higher during spiking than random moments (dashed line) across all timescales of decoding >2 ms. *Bottom* When mice occupied the new place field, the Pr(occupy new field | spks) increased after stimulation. Bars show Mann-Whitney U-test p < 0.01.

Next, we asked whether circuit modification persisted beyond track running. CA1 pyramidal cells that were more strongly recruited into sharp wave ripples (SWR)^21^ recorded during immobility on the track (N = 703 neurons, threshold for ripple recruitment = 2.5Hz, derived from median split of waking ripple activity for Stim and Control session) showed an increase in firing during post-RUN SWR recorded in the homecage, relative to pre-RUN session (Wilcoxon signed-rank test, p = 8.5^−8^). This increase was greater than that observed for pyramidal cells (N =672 neurons) that were less recruited into waking SWR (Wilcoxon signed-rank test, p = 1.69^−9^; Figure 3A), consistent with the well-known phenomenon of ‘replay’ ^22,23^. Similarly, interneurons that fired more during waking SWR (N = 127) also showed a post-RUN versus pre-RUN gain in homecage SWR firing (Wilcoxon signed-rank test, p 1.94^−4^), compared to interneurons which were only weakly recruited during waking SWR (Figure 3B; N = 119; Mann-Whitney U-test, p = 0.003). Pyramidal cells that were optogenetically stimulated (N = 230) versus those that were not (N = 220) did not differ in their post-RUN versus pre-RUN gains (Figure 3C; Mann-Whitney U-test, p = 0.46), though marginally more than those (N = 291) recorded in Control sessions (Figure 3C; p = 0.02). Thus, unlike the enhancement in ripple recruitment observed for those neurons that were spontaneously active during waking SPWs, optogenetic stimulation did not enhance post-RUN ripple-related activity, relative to pre-RUN recordings. In contrast, interneurons driven synaptically on the stimulated shank showed a large increase in SWR recruitment as compared to those recorded on the non-stimulated shank (Mann-Whitney U-test, p = 0.005) and compared to Control sessions (Mann-Whitney U-test, p = 0.005).

**Figure 3.**
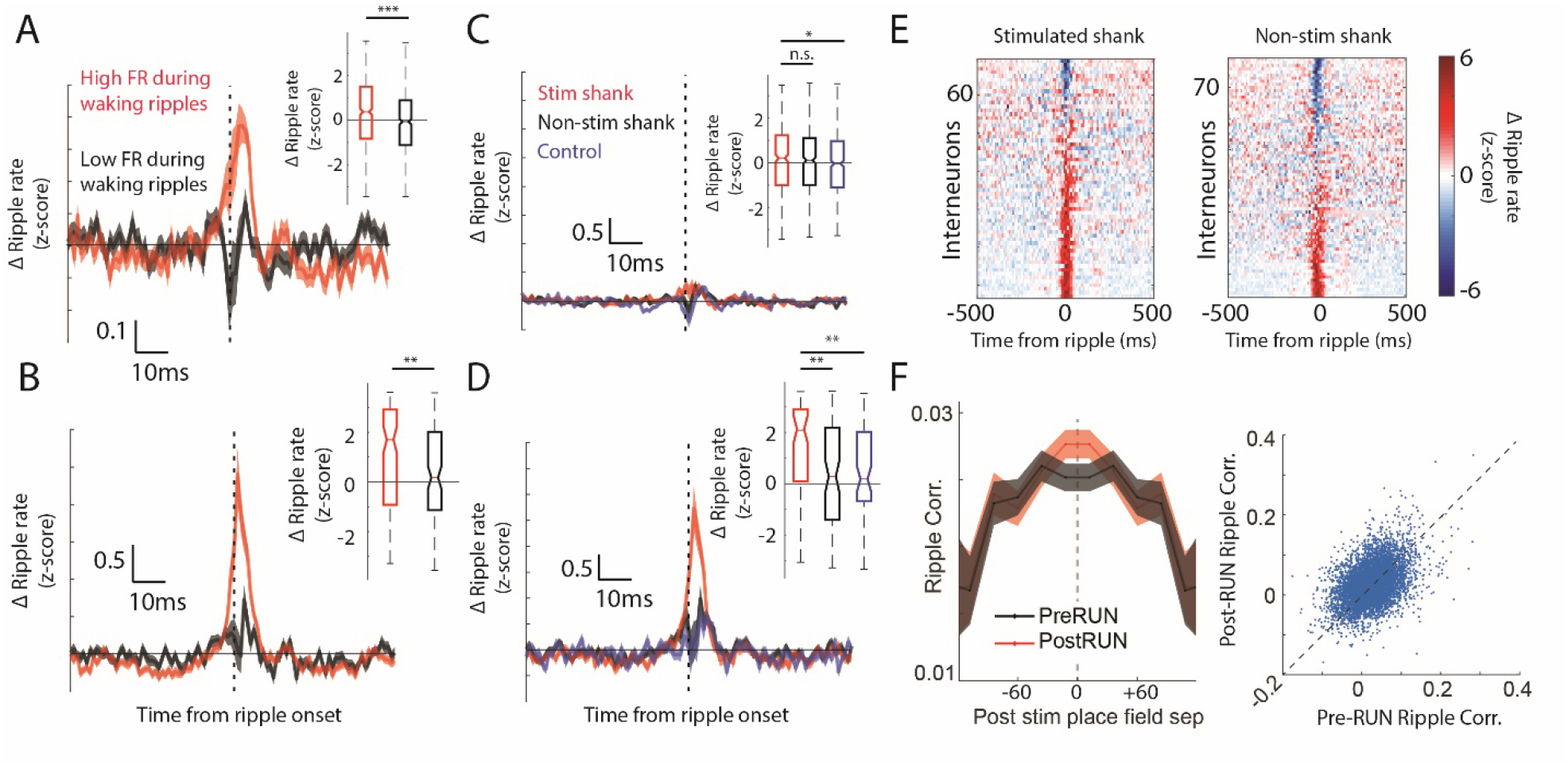
Induced place fields affect neural recruitment and membership in ripple events. *(A)* Pyramidal cells that fired during waking ripples on the track (red = upper 50%) showed greater gain in firing (post-RUN - pre-RUN) during ripples recorded in the homecage (median ± SEM) compared to those that participated less during waking ripples (black = lower 50%). *Inset:* median rate changes during ripples. *(B)* Interneurons recruited into ripples on the track also increased recruitment into homecage ripples. *(C)* Neurons recorded on the stimulated shank (red) were not recruited into subsequent ripples any more than simultaneously recorded non-stimulated neurons (black), or neurons recorded in Control session (blue). *(D,E)* Interneurons recorded on the stimulated shank show increased ripple recruitment. *(F) Left*, The distance between post-stimulation place fields is correlated with the pairwise correlation in ripple activity for SWRs recorded both prior to, and after, track running. *Right*, The pre-RUN pairwise ripple co-modulation is strongly correlated (Pearson r = 0.42, p = 2.1^−265^) with the post-RUN ripple co-modulation. *** p < .001, ** p <.01, * p < .05.

To test how stimulation changed the pattern of neural co-activity during ripples, the mean firing rate per ripple was correlated for each pair of neurons before and after track running. We focused on pairs in which one or both neurons shifted place fields after stimulations (N = 6,405 pairs). The degree of co-fluctuation (Spearman R) during ripples prior to track running strongly correlated with co-fluctuations after stimulation (Figure 3F; Post-RUN vs Pre-RUN co-fluctuation, Pearson corr. = 0.42, p = 2.1^−265^). To factor out this significant baseline stability, multiple regression analysis was used to measure how the distance between post-stimulation place fields predicted post-RUN ripple coupling, accounting for this pre-RUN baseline. The absolute distance between place fields post-stimulation negatively correlated with co-activity during post-RUN ripples (Figure 3F; t-stat = -3.08, p = 0.002), showing enhanced coupling for neurons whose place fields were nearby after optogenetically-induced remapping^22^. This relationship held even when using the pre-stimulation place field separation as an additional regressor (t-stat = -2.76, p = 0.006). The co-fluctuation of activity during pre-RUN ripples also negatively correlated with the distances between post-stimulation place fields (Figure 3F; t-stat = -3.26, p = 0.001), even after regressing out the distance between place fields prior to stimulation (t-stat = -2.7, p = 0.007), consistent with the observation that the order of place fields in a new environment is “preplayed” prior to that novel experience^11^.

We hypothesized that the changes in SWR participation of synaptically activated interneurons and the place field remapping, particularly of non-stimulated cells, was due to a reorganization of lateral inhibition. To explore this hypothesis, we examined spike transmission between pairs of monosynaptically connected pyramidal cells and interneurons (PYR-INT)^18,24^. To track how spike transmission changed over time, we developed a generalized linear model (see Supplemental Methods) to measure changes in the influence of the presynaptic drive to the postsynaptic interneuron while regressing out changes in postsynaptic firing rate (Figure 4A; Figure S4). We found that putative synaptic coupling strength, as approximated by our spike transmission measure, varied 64.7±0.5% around the mean over the recording session (Figure 4A,B, Figure S4). These temporal fluctuations were independent of the presynaptic rate (Spearman R = 0.0124, p = 0.077, N = 1,771 pairs), and correlated with the postsynaptic rate (R = 0.38, p < 0.001, N = 1,771 pairs), likely reflecting an effect of postsynaptic excitability (Figure S5A-B). Yet, spike transmission between neuron pairs sharing the same postsynaptic interneuron was only moderately correlated with each other (R = 0.24, p < 0.001, N = 12,581 convergent pairs), suggesting that the temporal dynamics of PYR-INT spike transmission cannot be fully explained by firing rate fluctuations alone (Figure S5C).

**Figure 4.**
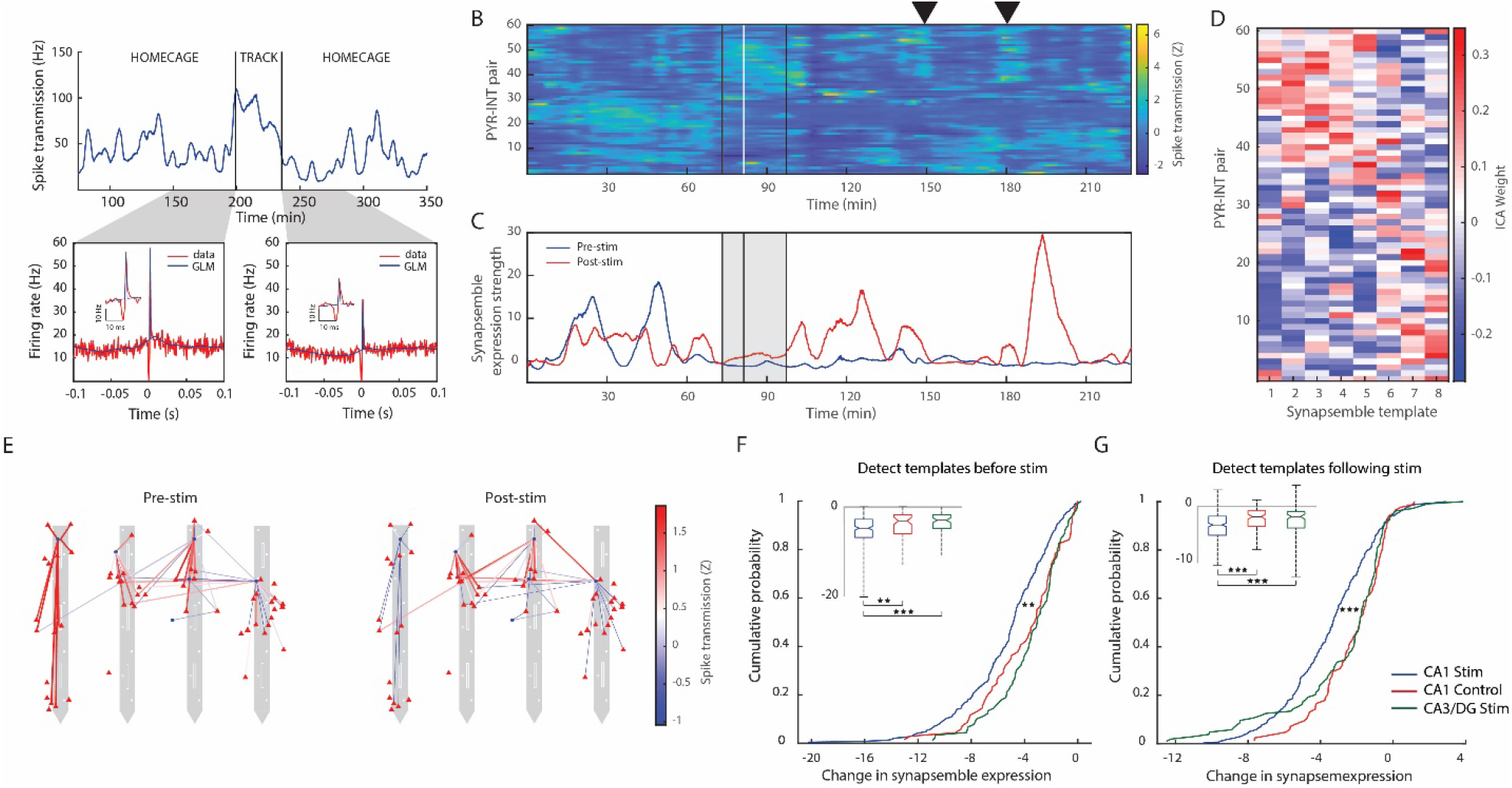
Optogenetic stimulation reorganized spike transmission between pyramidal cells and interneurons in CA1, but not CA3 or DG. *(A) Top*, Spike transmission (in Hz) between an example PYR-INT pair reflecting a GLM-based estimate of the increase in postsynaptic rate due to the presynaptic spikes (see Supplementary methods). *Bottom*, Observed and GLM-predicted cross-correlograms for the example PYR-INT pair, computed in 50-minute home cage periods before and after track running (see Supplementary methods). *Insets* highlight the peak at a finer time scale. *(B)* Z-scored spike transmission time series between 61 PYR-INT pairs in a single session. Black vertical lines: start and end of track running. White line, first optogenetic stimulation trial. ▴ highlights a recurring spike transmission ensemble. *(C)* Time-resolved expression strengths of two PYR-INT spike transmission ensembles (‘synapsembles’, see Supplementary methods) detected during Pre-stim (blue) and Post-stim (red) epochs. *(D)* Templates of PYR-INT ensembles detected prior to optogenetic stimulation. *(E)* Mean Z-scored spike transmission maps during periods of high ensemble expression (>10) for Pre-stim and Post-stim synapsembles shown in Panel C. Pyramidal cells (red triangles) and interneurons (blue circles) are superimposed on the recording sites (white squares) of a four-shank µLED probe. *(F, G)* Stimulation induced larger changes in CA1 synapsemble expression (blue, N = 243) than in CA1 control sessions (red, N = 69; p < 0.01; Mann-Whitney U-test) and in combined CA3/DG stimulation sessions (green, N = 90; p < 0.001). Synapsembles were detected prior to (*F)* or after *(G)* optogenetic stimulation; the halfway point was used in CA1 Control sessions. *Insets* show the same data as whisker plots.

Spike transmission between simultaneously recorded PYR-INT pairs tended to co-fluctuate, with pairs that increased and decreased at common moments (Figure 4B). To quantify how spike transmission across PYR-INT pairs is coordinated and changes as a result of optogenetic stimulation, we extracted patterns of significant coactivity using a combination of PCA and ICA (see Supplemental Methods) to define a vector of weights for the contribution of each PYR-INT pair to each synaptic state (Figure S4), referred to here as ‘synapsembles’^25^ (Figure 4 B-E). Each synapsemble was detected prior to, or after, stimulation, and its expression strength was tracked throughout the session (see Supplementary Methods). The synapsembles defined by pre-stimulation activity were re-expressed more weakly after optogenetic perturbation compared to those detected in CA1 Control sessions (p < 0.01, Mann-Whitney U-test) and in CA3/DG Stim sessions (p < 0.001; Figure 4F). These results also held when controlling for differences in baseline synapsemble expression strengths between optogenetic perturbation and Control conditions (Figure S5D). Identical results were obtained when synapsembles were identified *following* optogenetic perturbation and compared to their expression prior to perturbation (Figure 4G; Figure S5E). Since place field remapping and changes in synapsemble expression were observed in CA1, and not in CA3 and the dentate gyrus (Figure 2), we hypothesize that rearrangement of synaptic strengths between pyramidal cells and interneurons contributed to place field remapping.

In contrast to previous studies that manipulated and monitored single cells ^1,14,15^, we observed remapping both at, and away from, the stimulation site and in both stimulated and non-stimulated neurons. New place fields tended to emerge in places where there was weak prior drive ^10,11,16^ and preexisting coupling with co-tuned neurons, an indication that preconfigured dynamics both predicted and constrained the emergence of new place fields ^26^. Moreover, most stimulated CA1 neurons remained stable, and neurons in CA3 and the dentate gyrus showed no significant remapping. We hypothesize that the changes observed in area CA1 were brought about by rapid redistribution of recurrent and lateral inhibition. This hypothesis is also supported by the higher level of place field plasticity in the CA1 region as opposed to the CA3/dentate region ^27-29^, regions with strong recurrent excitation^30^. Overall, our findings suggest that the encoding of novel information within hippocampal circuits is constrained and guided by a backbone of preexisting repertoire of states ^9-12,26^.

## Acknowledgements

We would like to thank Zachery Saccomano, Abed Ghanbari, and Loren Frank for providing software used in the present study and Yunchang Zhang and Sinan Kokuuslu for their help with data collection. We would like to thank Manuel Valero, Thomas Hainmueller, Antonio Fernández-Ruiz, and Shy Shohamy for useful feedback on the manuscript prior to publication. This work was supported by NIMH K99 MH118423, NIH MH54671, NIH MH107396, and NS 090583, NSF PIRE grant (#1545858), U19 NS107616 and U19 NS104590.

---

**Supplemental Figure 1.**
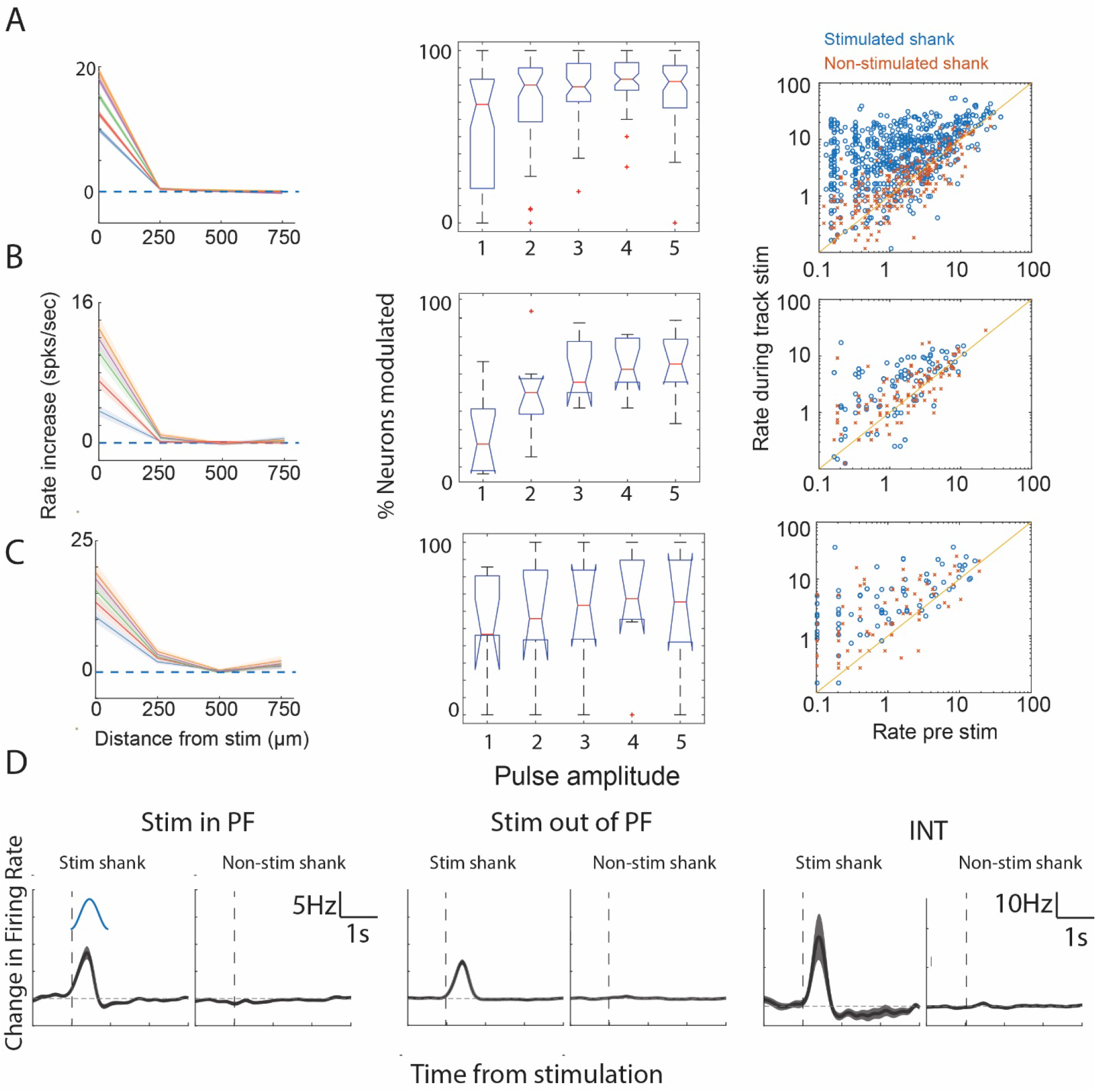
Focal optogenetic stimulation in CA1, CA3 and the dentate gyrus. *(A) Left*, In CA1, in the homecage, five calibration pulses (100ms) of different intensities were given such that the minimum reliably evoked firing and the maximum evoked a population oscillation. Strong increases in pyramidal cell firing was observed on the stimulated, and not neighboring shanks. *Middle*, The percentage of pyramidal cells stimulated on the illuminated shank during homecage stimulation. *Right*, The firing rates on the stimulated and non-stimulated shanks in the stimulated location on the trials before optogenetic drive compared to the rate during the optogenetic drive. *(B)* Same as A for principal cells in CA3. *(C)* Same as A for principal cells in the dentate gyrus. *(D)* Track stimulation for pyramidal cells in their existing place field and outside place field and interneurons in CA1.

**Supplemental Figure 2.**
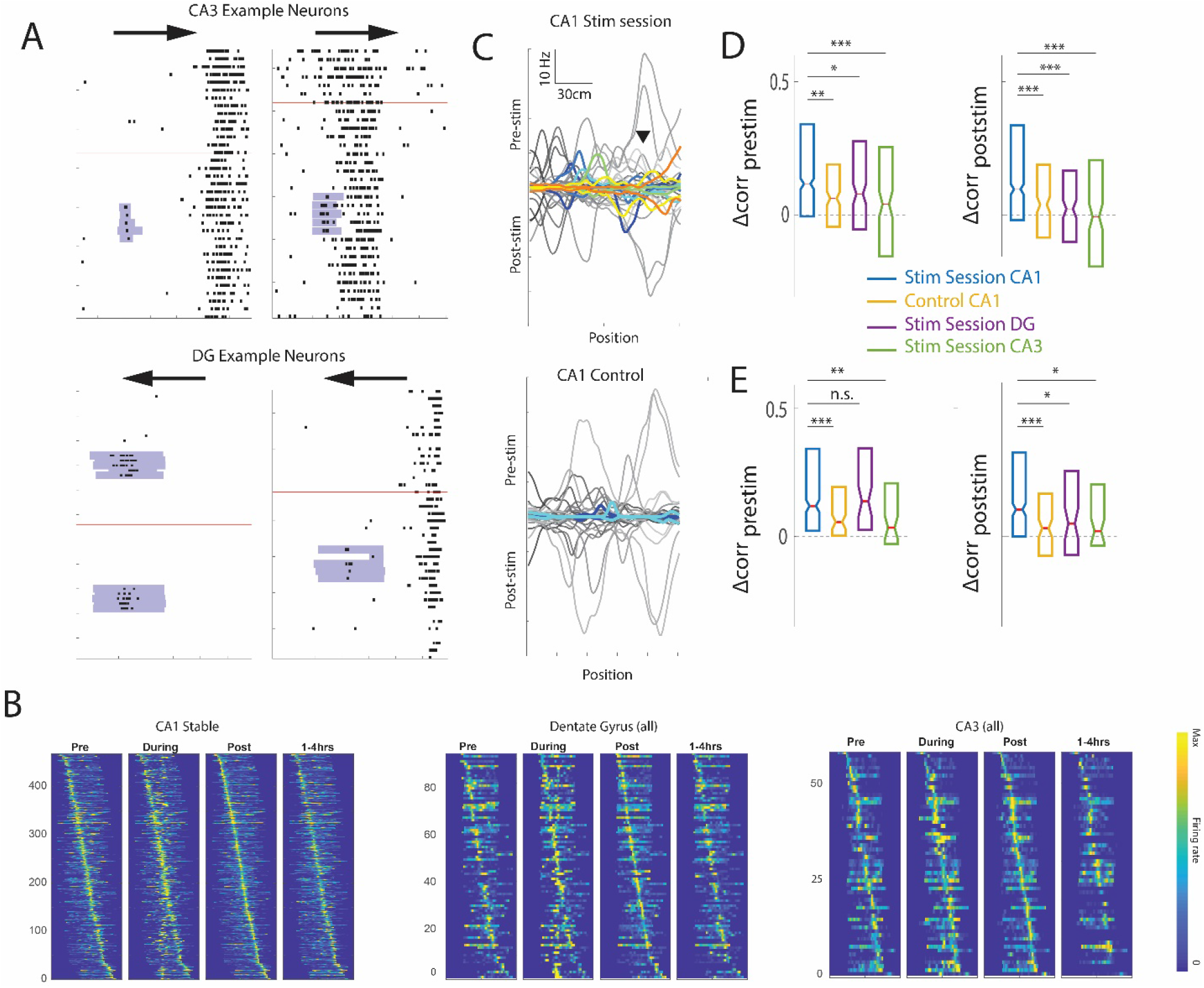
Optogenetically induced place field remapping is stronger in the CA1 region as compared to CA3 and dentate. *(A)* Example CA3 and dentate gyrus principal cells that maintained stable firing after optogenetic drive. *(B) Left*, All CA1 place fields that did not remap during CA1 stim sessions. *Middle*, All dentate gyrus place fields during stimulation sessions. *Right*, All CA3 place fields during stimulation sessions. *(C) Top*, remapping CA1 place fields in an example stimulation session as in Figure 1C. *Bottom*, Mostly stable (grey) place fields in a Control session. Remapping place fields are colored. *(D)* Larger pre vs post template mismatch for the non-stimulated running direction, Δcorr_prestim_ *(left)* and Δcorr_poststim_ *(right)*, for CA1 stimulation sessions as compared to Control sessions and CA3 and dentate gyrus stimulation session, see Figure 1F,G. *(E)* Same as Figure 1F,G using a spatial information (SI) criterion (SI_pre_ or SI_post_> 2.0 for place cell inclusion). *** p < .001, ** p <.01, * p < .05.

**Supplemental Figure 3.**
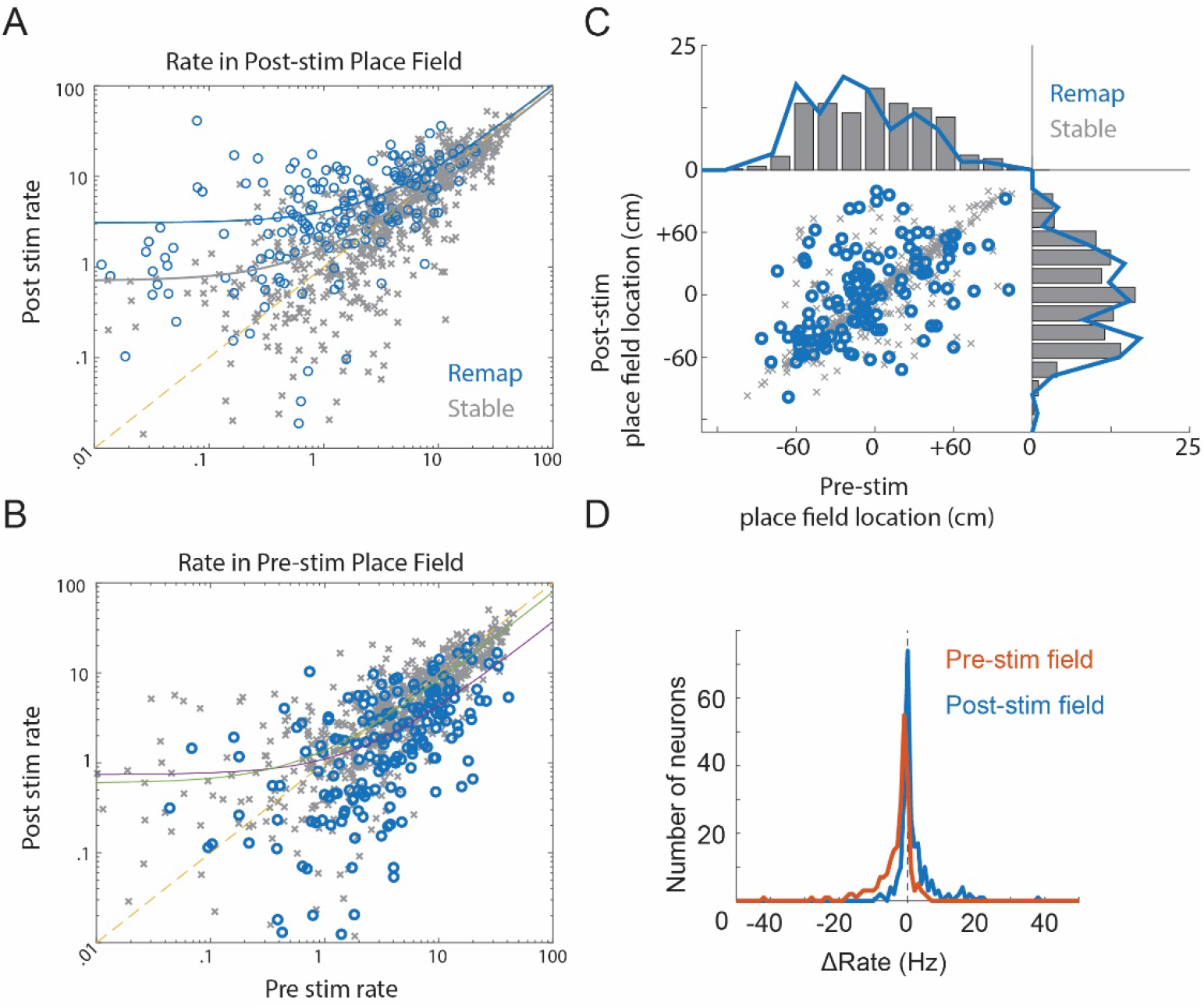
Remapping in the CA1 region occurred through increasing rates in randomly distributed locations on the track and decreasing rates in existing place field. *(A)* Though firing increased in the new place field (Δrate = 2.84 ± 0.34 Hz, signtest p = 8.36^−16^), there was also a strong correlation in the pre-vs post-stim rate correlation in the new field (Pearson r = 0.56, p = 8.4^−21^). To avoid regression to the mean, the location of the place field location was taken as the track position with the peak firing on odd trials; the reported firing rates at those locations were computed for even trials. The slope relating pre- and post-stimulation firing rate did not differ between remapping (slope = 0.96, CI: 0.80-1.11) and stable cells (slope = 0.87, CI: 0.84-0.90), though the y-intercept was higher (p < 0.0001) for remapping cells, as confirmed by a significant main effect (p = 7.5^−14^) and insignificant group x pre-stim rate interaction (p = 0.27) in an ANCOVA comparing remapping and stable cells. *(B)* After stimulation, firing rate decreased in the existing place field for remapping cells (Δrate = -2.2 ± 0.32 Hz, Mann-Whitney U-test p = 1.0^−7^). The slope relating pre- and post-stimulation firing rate significantly differed between remapping (slope = 036, CI: 0.33-0.39) and stable cells (slope = 0.78, CI: 0.76-0.81, p < 0.0001), as confirmed by a significant main effect (p < 1.0^−10^) and group x pre-stim rate interaction (p < 1.0^−10^) in an ANCOVA comparing remapping and stable cells, showing larger rate decreases for neurons with existing place fields. *(C)* On the population level, rate decrease in the old field was equal to the rate increase in the new fields (Mann-Whitney U-test p = 0.08), though for single cells, there was no correlation between the rate decrease in the old field and the rate increase in the new field (Pearson r = -0.02, p = 0.74) *(D)* There was no over-representation of the stimulated location (place fields < 20cm to stim location) post-stim (binomial test, p = 0.58), as new place fields could emerge or disappear anywhere on the track. *** p < .001, ** p <.01, * p < .05.

**Supplemental Figure 4.**
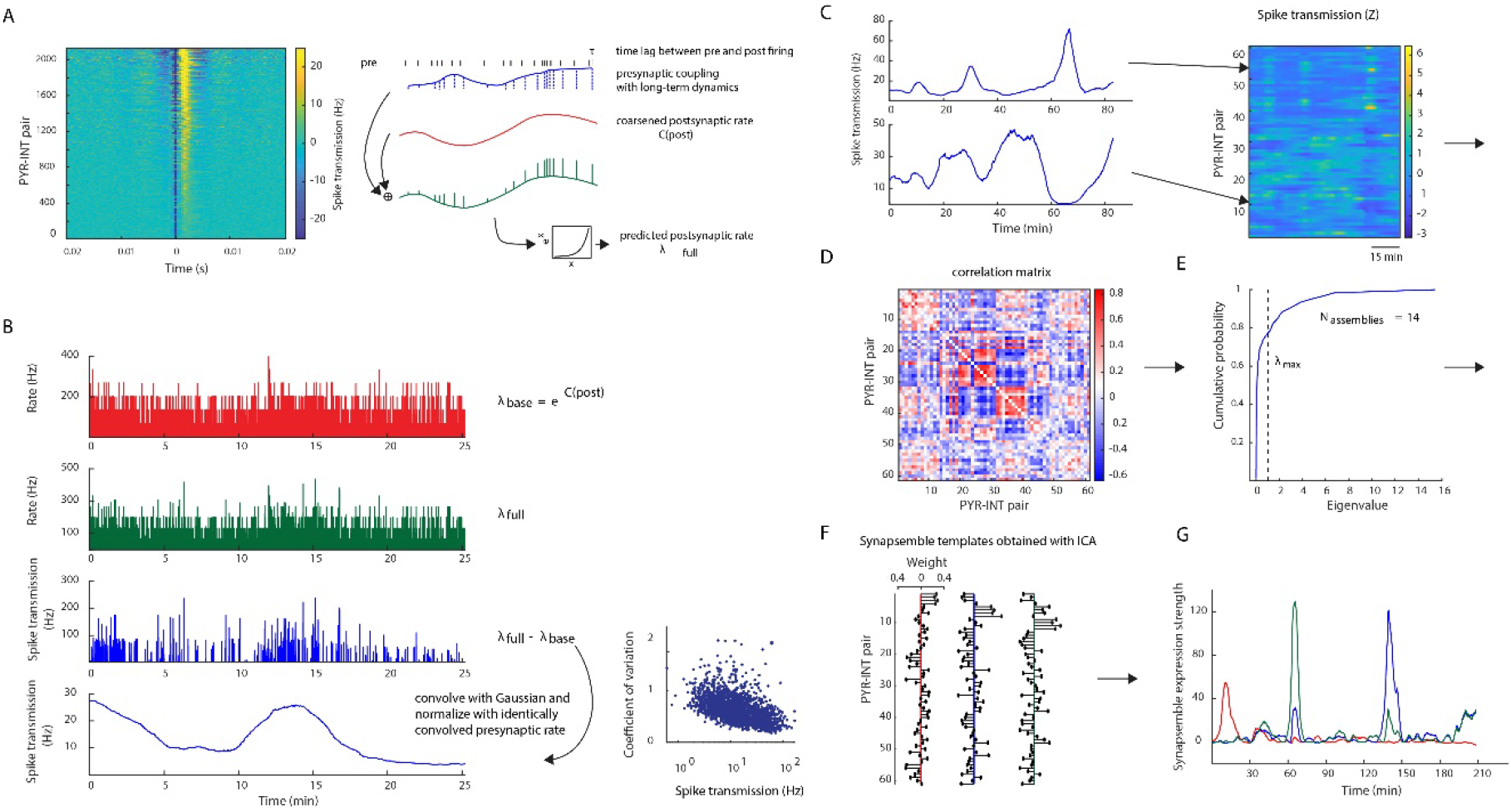
Calculation of time-resolved spike transmission and synapsemble detection. *(A) Left*, Monosynaptically connected pairs of pyramidal cells and interneurons (N = 2132) were detected as described in^18^. Baseline corrected cross-correlograms (CCGs) of all PYR-INT pairs are ordered by spike transmission strength, defined as excess rate in the 0.8–2.8 ms bins above the baseline^18^. *Right*, Schematic of a generalized linear model used to compute a time-resolved estimate of spike transmission. Before being passed through the spiking nonlinearity (green bottom trace), the postsynaptic rate of the interneuron is modeled as a linear combination of slow time varying (binned at 15ms and linearly interpolated) postsynaptic rate (red) and a time-varying rate-gain at the moment just after presynaptic spiking (blue). *(B) Left*, To estimate the time-resolved spike transmission between a PYR-INT pair, the coarsened postsynaptic rate (red, *λ*_*base*_) was subtracted from the full model (green, *λ*_*full*_) described in Panel A. This difference (blue, *λ*_*diff*_) reflects the time-varying postsynaptic rate at monosynaptic latency after each presynaptic spike, over and above what can be expected from slow changes in the postsynaptic rate alone. In order to obtain a smooth estimate of this quantity in units of rate per presynaptic spike, *λ*_*diff*_ was convolved with a Gaussian (SD = 120 seconds) and divided by the presynaptic spike train convolved in the same manner. *Right*, The magnitude of fluctuations in spike transmission probability was quantified using the coefficient of variation (CV = 0.647±0.005). *(C) Left*, Spike transmission time series of two PYR-INT pairs (left) are shown in an example session (61 pairs). *Right*, The time series are downsampled to 10 Hz, z-scored, and represented as a matrix. *(D)* Correlation matrix of all pairs in this session. Positive off-diagonal values reflect pairs whose spike transmissions co-fluctuate. *(E)* PCA was applied to the correlation matrix, and the number of significant spike transmission coactivation patterns (‘synapsembles’) was estimated as the number of eigenvalues exceeding the analytical threshold *λ*_*max*_ based on the Marcenko-Pastur distribution. *(F)* Z-scored data (as shown in Panel C) was projected onto the subspace of PCs whose eigenvalues exceeded *λ*_*max*_, and ICA was used to identify synapsemble templates. Example synapsembles are represented as a whicker plot. *(G)* Time-resolved expression strengths of the 3 synapsembles shown in Panel F. The synapsemble expression strength is defined as the bin-by-bin squared projection of the z-scored excess synchronies onto a given synapsemble template (see Supplementary Methods).

**Supplemental Figure 5.**
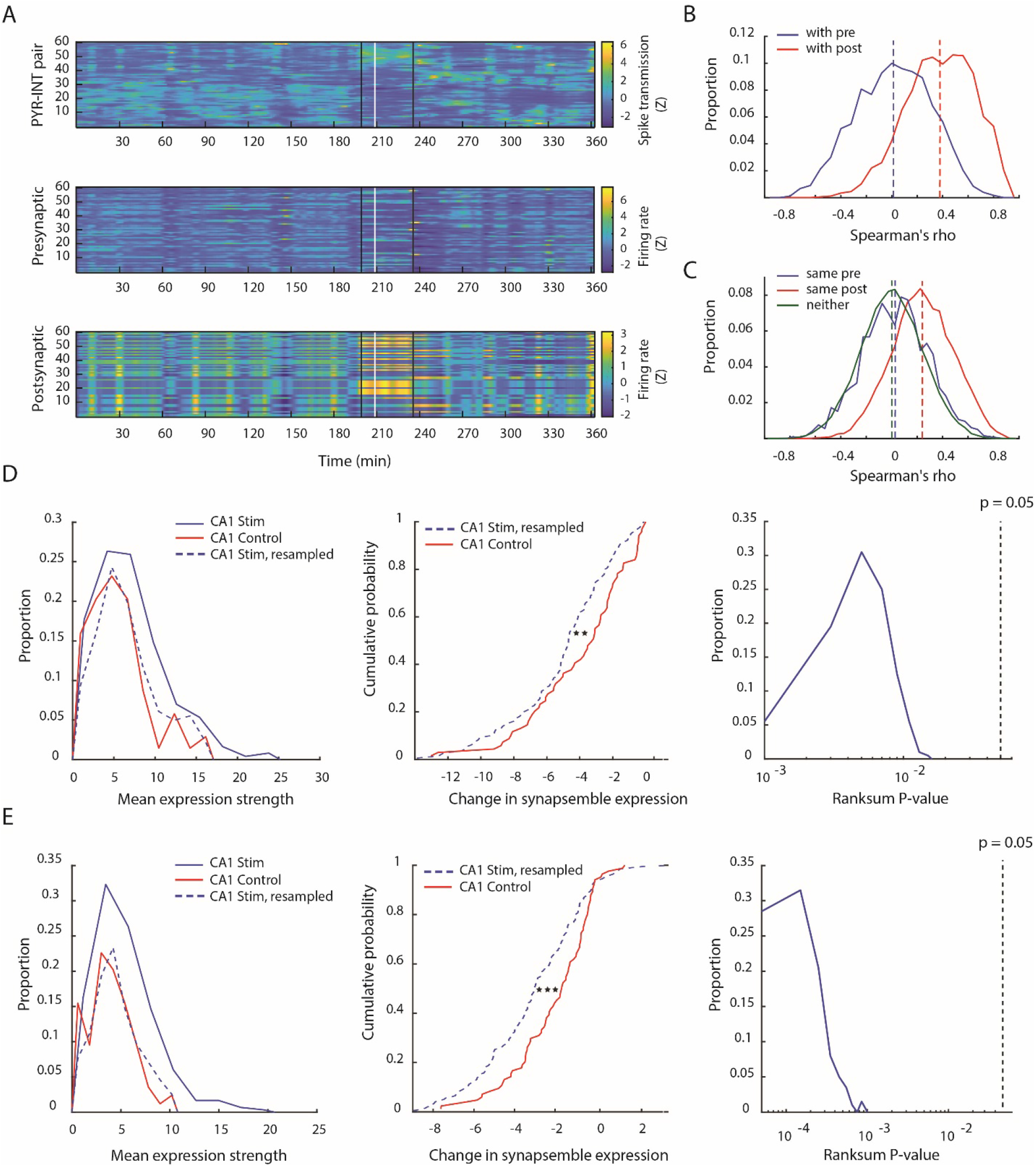
Spike transmission is not fully accounted for by rate, and synapsemble rearrangements are robust with respect to differences in baseline expression. *(A) Top* Spike transmission time series in a session different from that shown in Figure 4B. *Middle* Z-scored firing rates of the presynaptic pyramidal cells (ordered according to pair membership above). Spikes were convolved in the same way as the baseline corrected spike transmission time series (*λ*_*diff*_, see Supplementary methods). *Bottom*, Same as middle, but for postsynaptic interneurons. *(B)* Distributions of Spearman correlation coefficients between all presynaptic PYR rates (blue, N = 1,771, R = 0.0124) and spike transmission time series of the pairs they belong to and between postsynaptic INT rates (red, N = 1,771, R = 0.38) and the spike transmission time series of the pairs they belong to. Note that firing rate co-variation of the presynaptic pyramidal neurons does not affect spike transmission measure. *(C)* Spearman correlation coefficient distribution among excess synchrony time series sharing a presynaptic PYR (blue, N = 1,977, R = 0.027), a postsynaptic INT (red, N = 12,581, R = 0.24), or sharing neither (green, N = 42,859, R = 0.0003). *(D) Left*, Distributions of mean synapsemble expression strengths during Pre-stim recording (when synapsembles were detected) were different between CA1 stim (blue, N = 243) and control (red, N = 69) sessions (p > 0.05, Mann-Whitney U-test) samples. We considered that changes in synapsemble expressions (Figure 4F,G) could be a statistical artifact due to this difference in baseline. To control for this possibility, we sampled from the CA1 stim distribution according to the empirical cumulative distribution function of the CA1 control distribution. This yielded a surrogate CA1 stim distribution (blue dashed) that was no different from CA1 control (p < 0.01, Mann-Whitney U-test). *Middle*, Distributions of changes in mean synapsemble expression strengths between CA1 Control and resampled CA1 stim were significantly different (p < 0.01, Mann-Whitney U-test), supporting the result in Figure 4F. *Right*, The distribution of Mann-Whitney U-test p-values for N = 200 independently resampled CA1 stim distributions lies below the significance threshold (p = 0.05), suggesting that the results of this control analysis are robust with respect to the randomness of resampling. *(E)* Same as D), except synapsembles were detected during Post-stim recording.

## Supplemental Methods

### Subjects

Homozygous T29-1. CaMKIIα-Cre line T29-1 transgenic mice (Jackson Laboratory #005359) were crossed with homozygous Ai32 mice (Jackson Laboratory #012569) to express channelrhodopsin2 (ChR2) in neurons expressing male and female CaMKIIα in F1 hybrid mice (N = 6; 4 male; 25-40g, 30-50 weeks of age). After implantation, animals were housed individually on a reversed 12/12 h day/night schedule. Following one week of recovery, mice were recorded 5-7 days/week for two months before being euthanized with pentobarbital cocktail (Euthasol^®^, transcardial 300 mg/kg) and perfused with formalin (10%). All experiments were conducted in accordance with the Institutional Animal Care and Use Committee of New York University Medical Center.

### Task

Mice were trained to run laps on a linear track (120 cm long, 4cm wide) to retrieve water reward (5-10µL) at each end. Before implantation, water access was restricted and was only available as reward on a linear track and ad libitum for 30 minutes at the end of each day. After mice reliably ran 50 trials in under an hour (∼1week daily training), free water access was restored for at least two days, and surgery was scheduled.

For a typical recording session, a one hour baseline recording was conducted in the mouse’s home cage, followed by a calibration of light intensity for optogenetic stimulation. Then mice ran 25-30 trials in a morning session, followed by 2-3 hours of homecage recording and another 25 trials in an afternoon/evening session, followed by redelivery of the calibrating light pulses in the homecage and ad libitum water access. On Control sessions, the calibration pulses were given though no stimulation was delivered on the track.

To deliver optogenetic stimulation at a fixed location for 1 s, an infrared sensor was placed at a random location on the track. Sensors were also placed at each end to control water delivery. An Arduino detected beam breaks to activate solenoids to deliver water and to send a TTL pulse to a DAQ (CED Power 1401 Cambridge, UK) which delivered voltage control signals to the integrated µLEDs.

### Surgery

Mice were anesthetized with 1.5-2% isoflurane (2 L/min) and provided with a local anesthetic to the incision site (bupivicaine at .05 mg/kg, 2.5 mg/ml, S.C.). The skull was cleaned with saline and hydrogen peroxide and ground wires (bare stainless steel) were positioned intracranially over the cerebellum. The skull was then coated with Optibond (Kerr Dental, Brea, CA) and a craniotomy (∼1.5 × 0.5 mm) was performed at AP -2.2, ML -2.0 (left hemisphere), 45°angle from the midline. The dura was removed and the probe was implanted ∼0.5mm into the cortex. The probe and custom driver were cemented to the skull with C & B Metabond Quick Adhesive Cement (Parkell) and Unifast Trad acrylic (GC America). The craniotomy was capped with a mixture of mineral oil (one part) and dental wax (three parts), and a Faraday cage was constructed using copper mesh and connected to the cerebellar ground wire. Following surgery, an opioid analgesic was injected (Buprenex at 0.06 mg/kg, 0.015 mg/ml, IM) and given as needed for the next 1-3 days.

### Recording and stimulation

Neural data was acquired using 32 site, 4-shank µLED probes^1^ (Neuralight, MI). Data were amplified and digitized at 30kHz with Intan amplifier boards (RHD2132/RHD2000 Evaluation System, Intan). µLEDs were controlled with voltage (2-3V generating 5-30µW of total light power) provided by a CED Power 1401 programmed with Spike2 (CED) which delivered light pulses (pre and post run) or 1s long sine waves (when the track IR beam was crossed). The animal’s position was monitored with a Basler camera (acA1300-60gmNIR, Graftek Imaging) sampling at 30Hz to detect head-mounted blue and red LEDs. Position was synchronized with neural data with TTLs signaling shutter position as well as a blinking LED (0.5 Hz) mounted 1m above the maze.

Blue light (centered emission at 460 nm) was delivered on one or two µLEDs (always 1 µLED/shank). To minimize artifact, the control voltage was held just under the forward voltage (2V). For calibrating light intensity, pulses (100ms, 1-1.5s inter-stimulus intervals) were delivered at 5 amplitudes (20 pulses/amplitude), where the maximum intensity in the homecage generated a population oscillation. This maximum level was used during track running, where the inhibitory tone is higher and equivalent stimulation evoked a weak response (Figure S1). On the track, 1s half sine waves were delivered when mice crossed an IR beam. Track stimulation only occurred in one running direction and, unless otherwise noted, was given for a block of 5 trials.

### Analysis

#### Unit isolation

Spikes were extracted and classified using Kilosort^2^. Global principal components were calculated (three per channel,8 channels/shank) and spikes were extracted from the highpass filtered wideband signal (3^rd^ order butterworth filter, passband: 0.5 – 15kHz). Manual unit curation was done using Klusters. Spike sorting quality was assessed with L-Ratio^3^, Isolation distance^3^, inter-spike interval violation, and visual inspection of cross-correlations suggestive of erroneous splitting of single units.

#### Cell type classification

Spike waveform (width and asymmetry), autocorrelation properties, and mean firing rate (mean inter-spike interval) were used to classify neurons into excitatory cells and interneurons. The autocorrelation was parameterized with a double exponential model:

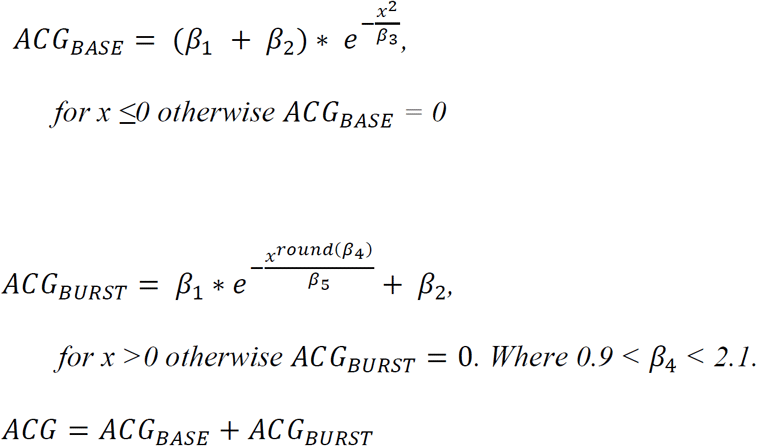

Units were defined by: rate, autocorrelation peak above baseline β_1_, spike width, and spike asymmetry. Then k-means clustering (k=2) was performed on the z-score normalized feature matrix which separated excitatory cells from putative interneurons (including fast spiking and regular spiking interneurons). The validity of the cluster labels was confirmed through the cross-correlation (CCG) analysis, revealing increased synchrony at synaptic time-scales (see *Synapsemble Analysis*).

#### Ripple detection

The local field potential (LFP) was extracted by low-pass filtering the 30kHz raw data (sinc filter with a 450 Hz cut-off band) and then downsampling to 1250Hz. For ripple detection, the LFP was bandpass-filtered (3^rd^ order Butterworth, passband: 130-200Hz), squared, and z-score normalized. Events with peak power > 5 standard deviation (SD), sustained power > 2 SD, and duration between 30-200 ms were detected. When available, such events were also detected on a non-hippocampal, ‘noise’ channels and events common to both (e.g., EMG artifacts) were excluded. Stimulation periods were also excluded. Ripple onset was the first moment when the bandpassed signal increased >2 st. devs.

#### Ripple co-fluctuation Analysis

Ripple start and stop times were taken as the moments when ripple-band power rose above and fell below 2 st. devs. of baseline. Rate per ripple was simply the number of spikes per neuron divided by the duration of the ripple event. Pairwise co-fluctuations were quantified through analysis of the Spearman correlation of ripple rates pre-(*Corr*_*preRUN*_) and postRUN(*Corr*_*postRUN*_. Multiple regression analyses were used to predict:

Post-RUN ripple correlations,

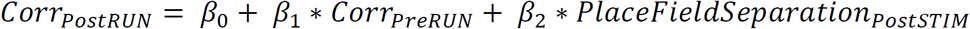

while accounting for pre-stimulation place fields

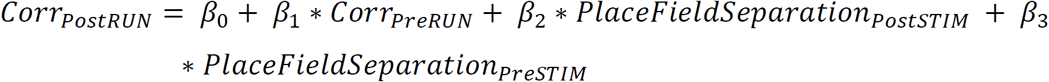

And preRUN ripple correlations

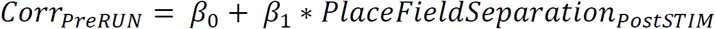

while accounting for pre-stimulation place fields

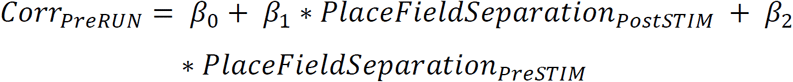

#### Place Field Remapping Analysis

The two-dimensional location of the mouse was linearized by projecting actual position onto the straight line of the track. A Kalman filter (2^nd^ order; locally quadratic) was used to derive a Bayesian MAP estimate of instantaneous speed and only moments with speed >1.5cm/sec were considered for place field analysis. Position was binned (100 bins, each 1.2cm) and the spike count and occupancy at each binned position was convolved with a Gaussian kernel (σ =5 spatial bins = 4.2 cm). Firing rate maps were calculated as the smoothed spike counts divided by the smoothed occupancy.

To calculate a trial-by-trial estimate of remapping, a template method was adopted. Two templates were calculated, one for place field maps prior to stimulation (premap) and another for post-stimulation place field maps (postmap). Both templates were correlated (Pearson’s correlation coefficient) with single trial rate maps (premap corr./postmap corr.), always excluding the correlated trial from the data from which the template was defined (i.e. the premap template is derived from data from trials 2-10 when correlated with trial 1). The mean difference between the template matches prior to stimulation, Δcorr_prestim gives a measure for predictiveness of poststimulation spatial tuning in generating the prestimulation rate maps. The mean difference between the template matches post stimulation, Δcorr_poststim provides the converse measure, the predictiveness of prestimulation place field tuning for post stimulation activity. To avoid firing rate thresholds, we opted to include any place field map that showed trial-to-trial spatial reliability (mean premap corr. > 0.25 for trials before stimulation and mean postmap corr. > 0.25 for trials after stimulation). Inclusion criteria using spatial information^4^ showed similar results (Figure S2). Inhibitory cells were excluded from place field analyses.

To further test how post-stimulation fields relate to pre-stimulation activity (Figure 2), we focused on neurons that showed reorganization of place field activity (Δcorr_poststim > 0.25 or Δcorr_prestim > 0.25) and neurons that showed sudden emergence of a new place field. The sudden emergence of a new field was detected using an adaptive filter where dynamic spatial tuning curves were estimated while regressing out changes in overall firing rate and rhythmicity ^5^. Fields were considered to emerge when the mean spatial intensity function jumped by at least 1.0 Hz and remained on average >1Hz.

#### Bayesian Decoding Analysis

We wished to test whether, prior to remapping, neurons are already part of an assembly tuned to the future place field location. To address this question, we hypothesized that when the remapping neuron fired, the other simultaneously recorded neurons should predict that the mouse occupies the future place field location. To quantify this prediction, we used Bayesian decoding to determine the posterior probability of occupying the new place field location (Pr(occupy new field | spks). Place field templates of pyramidal cells were calculated from post-stimulation trials using only odd trials. Bayesian decoding^6^ on ensemble spiking data was performed on pre-stimulation trials and even post-stimulation trials (even/odd cross-validation). The time window for calculating the instantaneous spike count observation was varied from 2-1000 ms in 20 logarithmically spaced bins (Figure 2C).

#### Synapsemble Analysis

A wealth of in-vitro data suggests that the strength of synaptic coupling (e.g., magnitude of the evoked PSP) changes following different pairing protocols. Our goal was to capture related changes in our dataset by estimating long-term changes in spike transmission between monosynaptically connected pyramidal cells and interneurons (detected as in^7^; Figure S4). In order to do this, we model the postsynaptic spike train using a generalized linear model (GLM) with the following two features: (1) a coarsened, slowly-evolving version of the postsynaptic rate (baseline term) and (2) a transient boost following the presynaptic spike whose magnitude varies with time (coupling term). The features are summed and passed through an exponential nonlinearity to yield the instantaneous postsynaptic rate *λ*_*post*_(*t*). The exponentiated coupling term can therefore be interpreted as a multiplicative, presynaptically-induced gain acting on an otherwise slowly evolving postsynaptic rate (the exponentiated baseline term). This separation of timescales assumption conveniently dodges the issue of capturing all parameters modulating the postsynaptic rate (e.g., theta, behavioral state). The conditional intensity function takes the following form:

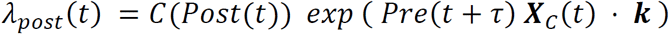

where *Pre* and *Post* are the pre- and postsynaptic spike trains binned at dt = 0.8 ms, ensuring that each bin has at most one spike. *τ* is the mode time lag between the pre- and postsynaptic spikes estimated from the raw CCG. C(Post) is the coarsened baseline rate obtained by counting postsynaptic spikes in 15 ms wide bins, expressing these as rates, and linearly interpolating at times corresponding to the centers of the dt-sized bins that were used to bin spikes. Note that in the absence of a presynaptic spike (i.e., *Pre*(*t* + *τ*) = 0), the predicted rate is equal to C(*Post*(*t*)). We model the slow changes in synaptic coupling ***X***_*C*_ · ***k*** with a linear combination of cubic B-splines with equally spaced knots ^8^. For each pair of neurons, the spacing of the knots (every 400-1000 seconds, in increments of 100) is selected by cross-validation using 100 second even/odd data splits. More specifically, the data are divided into 100 second data segments, such that odd segments are used for training, and even segments for testing. Because the parameters ***k*** are only constrained when *Pre*(*t* + *τ*) = 1, we fit the model in these bins exclusively. We use an LBFGS algorithm to minimize the convex negative log-likelihood with analytically computed gradients.

The instantaneous spike transmission – i.e., postsynaptic rate injected by a presynaptic spike – was estimated by taking the difference between the coarsened postsynaptic rate and that predicted by the full model:

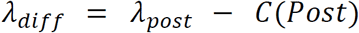

In order to obtain a smoothly varying estimate of spike transmission (i.e., even at times when the presynaptic neuron is not spiking), *λ*_*diff*_ was convolved using a Gaussian kernel with a standard deviation of 120 seconds. In order to get a bin-by-bin estimate of spike transmission (in Hz) per presynaptic spike, we normalized the resulting time series by the presynaptic spike train convolved in the same manner.

In order to find ensembles reflecting higher order coactivations among the spike transmission time series (i.e., synapsembles), we performed an unsupervised statistical analysis based on ICA ^9-11^. Briefly, spike transmission time series were represented as a matrix **z**, which was z-scored and downsampled to 10 Hz. The number of synapsembles in each session was based on the N principal components of the correlation matrix whose variances exceeded an analytical threshold based on the Marcenko-Pastur distribution describing variances expected for uncorrelated data. The high-dimensional activity matrix was projected onto the subspace spanned by these N principal components, and ICA was performed to extract synapsembles (each corresponds to an independent component). The expression strength of each synapsemble was computed as

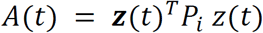

where *P*_*i*_ is the projection matrix (outer product, diagonal set to zero) of the i-th independent component. As such, A(t) quantifies the moment-to-moment similarity between an independent component (synapsemble template) and the instantaneous spike transmission pattern across all PYR-INT pairs.

In order to quantify synapsemble rearrangements across a relevant time point T, we defined the change in synapsemble expression as the difference between mean expression strength around T^11^. For stimulation sessions, T was the first stimulation event on the track, while for controls, we selected the halfway point on the track. In each case, differences in mean expression strengths were based on time intervals that included entire homecage periods flanking the track period. Synapsembles were detected either in the period before or following T. A negative value for the change in expression strength reflects that the synapsemble is reexpressed less prominently following T (if it was detected prior) or before T (if it was detected after).

**Supplemental Table 1.** Number of stimulations per session per region

|  | 0 stims (Control) | < 5 stims | 5 stims | >5 stims |
| --- | --- | --- | --- | --- |
| CA1 | 11 | 13 | 24 | 7 |
| CA3 | 0 | 5 | 7 | 4 |
| DG | 0 | 5 | 10 | 1 |

